# Spindle checkpoint silencing at kinetochores with submaximal microtubule occupancy

**DOI:** 10.1101/472407

**Authors:** Banafsheh Etemad, Abel Vertesy, Timo E.F. Kuijt, Carlos Sacristan, Alexander van Oudenaarden, Geert J.P.L. Kops

## Abstract

The spindle assembly checkpoint (SAC) ensures proper chromosome segregation by monitoring kinetochore-microtubule interactions. SAC proteins are shed from kinetochores once stable attachments are achieved. Human kinetochores consist of hundreds of SAC protein recruitment modules and bind up to 20 microtubules, raising the question how the SAC responds to intermediate attachment states. We show that the ‘RZZS-MAD1/2’ module of the SAC is removed from kinetochores at low microtubule occupancy and remains absent at higher occupancies, while the ‘BUB1/R1’ module is retained at substantial levels irrespective of attachment states. Artificially tuning the affinity of kinetochores for microtubules further shows that ~50% occupancy is sufficient to shed MAD2 and silence the SAC. Kinetochores thus responds as a single unit to shut down SAC signaling at submaximal occupancy states, but retains one SAC module. This may ensure continued SAC silencing on kinetochores with fluctuating occupancy states while maintaining the ability for fast SAC re-activation.

## Introduction

Errors in chromosome segregation cause aneuploid karyotypes, which are devastating to embryonic development and are strongly associated with cancer (de Wolf and Kops, 2017; Duijf et al., 2013; Hanahan and Weinberg, 2011; Ricke and van Deursen, 2013). To ensure proper chromosome segregation, the spindle assembly checkpoint (SAC) prevents anaphase initiation until all chromosomes are stably attached to spindle microtubules. These attachments are powered by kinetochores, specialized structures assembled on centromeric chromatin (Musacchio and Desai, 2017). Microtubule binding by kinetochores is mediated predominantly by the NDC80 complex (Cheeseman et al., 2006; DeLuca et al., 2002; DeLuca and Musacchio, 2012; Tooley and Stukenberg, 2011). When unbound by microtubules, however, this complex recruits the MPS1 kinase to kinetochores (Aravamudhan et al., 2015; Hiruma et al., 2015; Ji et al., 2015; Liu and Winey, 2012), where it initiates a cascade of events that culminates in production of the anaphase inhibitor. The cascade involves phosphorylation of the short linear MELT sequences in the kinetochore protein KNL1 to form the binding sites for the BUB3-bound SAC proteins BUBR1 and BUB1 (Krenn et al., 2014; Overlack et al., 2015; Primorac et al., 2013; Vleugel et al., 2013; Zhang et al., 2014). MPS1 also ensures localization of the MAD1-MAD2 complex, at least in part by promoting BUB1-MAD1 interactions (Kim et al., 2012; London and Biggins, 2014; Silió et al., 2015). MAD1-MAD2 recruitment additionally requires the RZZ (ROD-ZW10-Zwilch) kinetochore complex but the mechanism of this has not been elucidated (Caldas et al., 2015; Matson and Stukenberg, 2014; Silió et al., 2015). Although poorly understood at the molecular level, a subset of these SAC proteins then form a multiprotein assembly with potent anaphase inhibitory activity (Chao et al., 2012; Herzog et al., 2009; Kulukian et al., 2009; Sudakin et al., 2001).

Whereas recruitment of SAC proteins to kinetochores is essential for proper SAC activation, their removal is crucial for efficient SAC silencing and timely anaphase onset (Ballister et al., 2014; Ito et al., 2012; Jelluma et al., 2010; Kuijt et al., 2014; Maldonado and Kapoor, 2011). Microtubule attachments disrupt SAC signalling from kinetochores by mediating poleward transport of SAC proteins by the dynein motor complex (a process referred to as ‘stripping’) (Howell et al., 2001), and by affecting the balance of SAC-regulating kinases and phosphatases (Etemad and Kops, 2016; Funabiki and Wynne, 2013; Saurin, 2018). For example, RZZ-MAD1 is cargo of dynein via interactions with the kinetochore-specific dynactin adaptor Spindly (Barisic et al., 2010; Caldas et al., 2015; Chan et al., 2009; Gassmann et al., 2008; Kops et al., 2005; Silió et al., 2015). By contrast, BUB protein removal is dependent on inhibition of local MPS1 activity and reversal of MELT phosphorylations by the PP1 phosphatase (Etemad and Kops, 2016; Hiruma et al., 2015; Ji et al., 2015; London et al., 2012; Meadows et al., 2011; Nijenhuis et al., 2014; Rosenberg et al., 2011; Zhang et al., 2014).

The subcellular architecture of kinetochores is substantially more complex than illustrated above. A single human kinetochores contains ~240 NDC80 complexes likely configured in a lawn-like macro-structure (Suzuki et al., 2015; Zaytsev et al., 2014). This lawn can bind up to 20 microtubules that together form a so-called kinetochore (k-)fiber (DeLuca et al., 2005; McEwen et al., 2001; Nixon et al., 2015; Wendell et al., 1993). Likewise, when unbound by microtubules, a single human kinetochore binds hundreds of SAC modules (Howell et al., 2004; Vleugel et al., 2015). This subcellular complexity of kinetochores raises numerous questions about the response dynamics of SAC modules to increasing amounts of bound microtubules. Recent evidence points towards a model in which the SAC signal from kinetochores as a function of microtubule binding is not binary, but can exist in intermediate states: The total amount of MAD2 on kinetochores in cells correlates with the average mitotic delay imposed by the SAC (Collin et al., 2013), MAD1 removal is initiated before a full occupancy state is reached (Kuhn and Dumont, 2017), and a novel microtubule poison that reduces microtubule occupancy at kinetochores did not prevent SAC silencing (Dudka et al., 2018). It remains unclear though if SAC signalling is fully shut down only when kinetochores have acquired close to maximal microtubule occupancy, and if not, what occupancy state is sufficient for allowing mitotic exit (Burke and Stukenberg, 2008; Stukenberg and Burke, 2015). Furthermore, do all SAC modules behave similarly, or are there functionally relevant differences in the way they respond to different amounts of bound microtubules? We here address these questions by quantitative correlation imaging of SAC protein levels and microtubule occupancy at single kinetochores, and by assessing SAC activity and SAC protein amounts on kinetochores with experimentally manipulated average occupancy states.

## Results

### Simultaneous quantification of tubulin and kinetochore proteins on individual kinetochores

To examine the effect of intermediate states of kinetochore-microtubule attachment on SAC silencing, we wished to simultaneously quantify the relative amounts of SAC proteins and microtubules on individual kinetochores. To this end, cells were treated either with nocodazole to induce depolymerisation of microtubules, or with the proteasome inhibitor MG-132 to allow maturation of k-fibers without mitotic exit. Coldshock treatments were used to visualize k-fibers and remove other microtubules from the mitotic spindle (DeLuca, 2010; Polak et al., 2017). We stained cells for tubulin and CENP-C to mark kinetochores, and used line plot measurements on maximum or sum projection images to quantify their levels on individual kinetochores. Comparison of this method to other combinations of image processing and analysis showed similar results on the same images (**Sup. Figure 1A, B**). As expected, individual k-fibers in nocodazole-treated cells were absent, while those in MG-132-treated cells were of high intensity (**Sup. Figure 1C**). However, in agreement with previous reports (DeLuca et al., 2005; Maiato et al., 2006; Wendell et al., 1993), individual k-fibers in MG-132-treated metaphase cells displayed substantial intensity differences, indicative of non-uniform microtubule occupancies across metaphase kinetochores.

### Microtubule attachments evoke two types of responses on SAC proteins

To assess how the SAC modules responds to full occupancy on individual kinetochores, we next quantified levels of six SAC proteins and one SAC-regulating post-translation modification (KNL1-pT180, hereafter referred to as ‘pMELT’ (Vleugel et al., 2015)). As expected, SAC protein levels were high on kinetochores of nocodazole-treated cells that have no microtubule attachments (‘NULL’ condition), and were significantly reduced on those of cells treated with MG-132, which have acquired full microtubule occupancy (‘FULL’ condition) (Howell et al., 2004) (**Figure 1A, B**). We found substantial variance in the amount of accumulated SAC proteins on unattached kinetochores (**Figure 1B**). The high variance was also observed within single cells and independent of variance of more stable kinetochore components such as CENP-C (**Sup. Figure 2A, B**) or HEC1 (Collin et al., 2013). Of note, variability between kinetochores was independent on the method used for measuring local protein levels (**Sup. Figure 1A-C**), by differences between biological replicates (**Sup Figure 2C**), by different antibody penetration proficiencies (**Sup. Figure 2D, E**), or by kinetochore size (**Sup. Figure 2F**).

**Figure 1.**
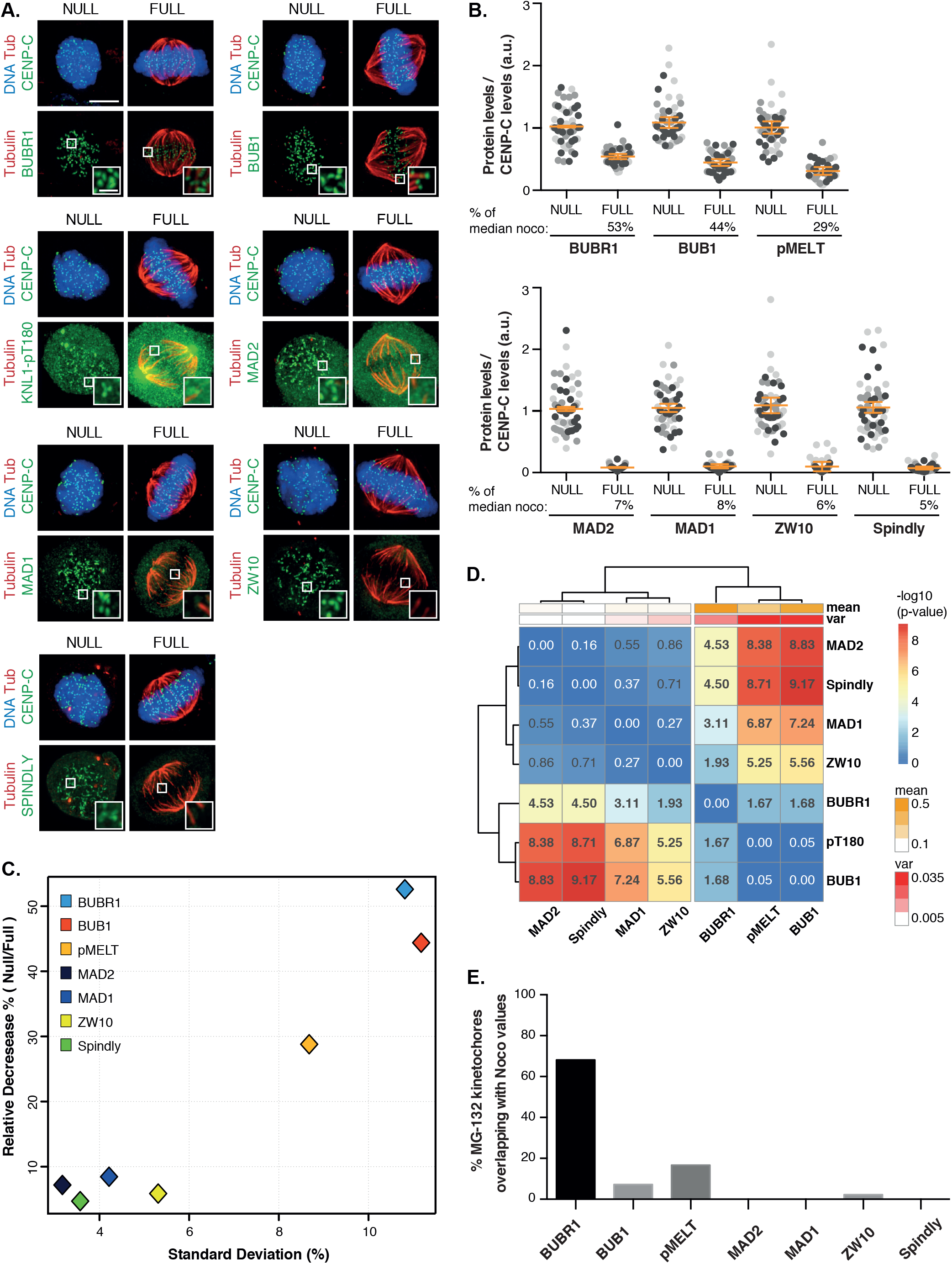
Microtubule attachments evoke two distinct SAC protein responses. **A-B)** Immunofluorescence images (A) and quantification of protein levels (B) on attached versus unattached kinetochores. Cells were synchronized and released in nocodazole (NULL) or treated with MG-132 (FULL) for two hours after mitotic entry. Each data point represents protein levels on a single kinetochore. Data from three independent experiments, represented by different shade of grey, are shown. Data is normalized to the median of the nocodazole levels measured in the same experiment to exclude batch effects. Error bars depict average of three experiments and SEM. At least 15 kinetochores in five cells or more were measured for each condition in every experiment. Channel colors of merged images match those of the labels. Scale bar, 5 *μ*m. Scale bar zoom-ins, 1 *μ*m. **C)** Plot depicting correlation between the relative protein decrease from NULL to FULL conditions and the standard deviation measured at FULL attachment (shown in B). **D)** Clustering heat map of P-values from pairwise Levene’s tests on the levels of proteins measured in the FULL condition. −log10(p-values) are displayed. Significant differences are in bold. Top bars represent variance (red) and average levels (orange) of each protein. **E)** Bar graph showing the fraction of FULL kinetochores that have similar levels of SAC protein as their NULL counterparts. MAD2, MAD1, and Spindly have no overlap.

Although all imaged SAC proteins were substantially delocalized from kinetochores of metaphase cells, they showed non-uniform behaviour. MAD2, MAD1, Spindly and ZW10 levels decreased to below detection limit. In contrast, BUBR1, BUB1 and pMELT were maintained to detectable levels at fully attached kinetochores (29-53% of the median of the levels measured on unattached kinetochores), in agreement with previous observations (Ballister et al., 2014; Bomont et al., 2005; Howell et al., 2004; Martinez-Exposito et al., 1999; Skoufias et al., 2001) (**Figure 1A, B**). Moreover, considerable variation between kinetochores was observed for these proteins on fully attached kinetochores (**Figure 1C**, x-axis) and pairwise comparison of the variance of all SAC proteins resulted in clustering of high and low variance proteins into two distinct groups (**Figure 1D**). To ensure that the difference in measured variation was not technical, we calculated signal-to-noise ratios and found that they are similar for high- and low-variance proteins, supporting a biological origin of this pattern (**Sup. Figure 3**). In addition, a fraction of attached kinetochores had accumulated as much BUBR1, BUB1 and pMELT as some of their unattached counterparts (Auckland et al., 2017), which was never observed for MAD2, MAD1 and Spindly, and only to limited extent for ZW10 (**Figure 1E**). The results from quantitative immuno-imaging were verified using genome-edited cell lines that express N-terminal HA-mCherry-tagged versions of MAD2 or BUB1 from their endogenous locus, excluding differences between antibodies as a cause for differences between SAC protein behavior at kinetochores (**Sup. Figure 4**).

### SAC proteins respond to intermediate attachments states

To examine how SAC signalling from kinetochores responds to intermediate kinetochore-microtubule attachment states, we measured SAC protein and tubulin signal intensities on individual kinetochores with immature k-fibers. To this end, cells were fixed in prometaphase after release from a G2/M-boundary block (see M&M for details). The resulting population of kinetochores had a mixture of attachment states (**Sup. Figure 5A, B**), including unattached and fully attached, as evident from comparisons to simultaneous imaging of kinetochores from the ‘NULL’ and ‘FULL’ conditions (**Figure 2A-H**). We found that all SAC proteins were substantially reduced on kinetochores with ~30% or more microtubule occupancy (relative to average tubulin intensity in the ‘FULL’ condition). However, whereas most kinetochores had no or barely detectable MAD1, MAD2, ZW10 and Spindly (‘RZZS-MAD1/2’ group) at ~50% of average max occupancy, members of the ‘BUB1/R1’ group (KNL1-pT180, BUBR1, BUB1) remained clearly detectable and with substantial variability. Segmented linear regression and hierarchical clustering of various extracted features (**Sup. Figure 5B, C**) verified that occupancy response-curves separated into two clusters: one containing the profiles of MAD1, MAD2, Spindly and ZW10 and the other containing those of BUB1, BUBR1 and KNL1-pT180 (**Figure 2I**). This was consistent with the behaviour of these proteins to full attachment (**Figure 1A, B**) and suggests a mechanistic difference in their response to microtubule attachments. The clustering analysis predicted that members of the RZZS-MAD1/2 module should behave differently to identical occupancy than members of BUB1/R1 module when measured on the same kinetochore. Indeed, (partly) attached kinetochores with no detectable mCherry-MAD2 had a variety of BUB1 levels, and BUB1 displayed greater variability than MAD2 (**Figure 2J**). In contrast, the signal intensities of BUB1 and BUBR1 on the same kinetochores of prometaphase cells were strongly correlated (**Figure 2K**).

**Figure 2.**
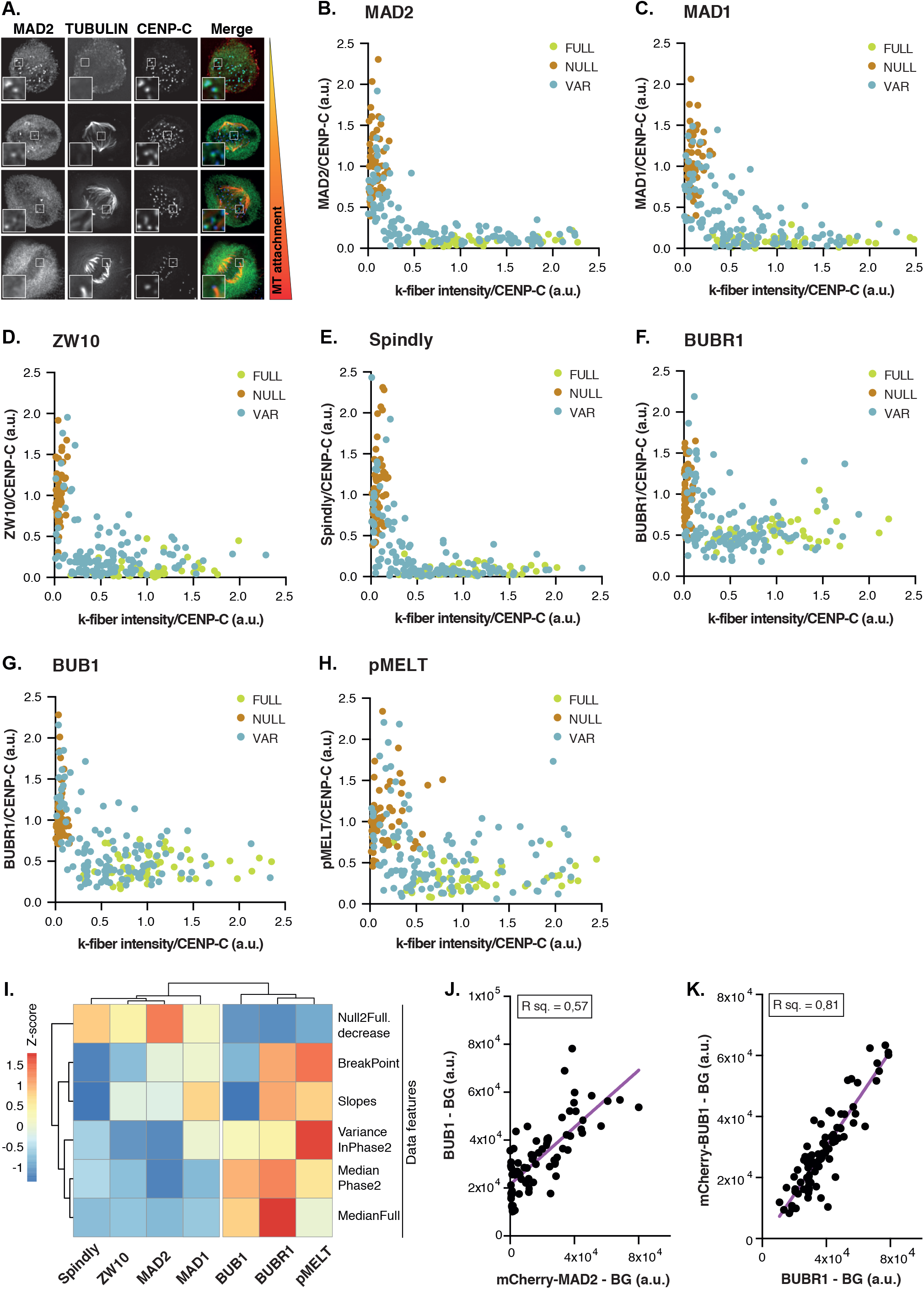
SAC proteins respond to intermediate attachments states. **A-H)** Representative images (A) and quantifications of levels of SAC proteins on kinetochores with different levels of intermediate attachments. Graphs (B-H) show data of three biological replicates. SAC protein levels of each experiment are normalized to the median levels measured in their respective NULL conditions, and tubulin levels are normalized to the median levels measured in the FULL condition. For each protein, at least 15 kinetochores in >5 cells are measured per experiments in the FULL and NULL conditions. For the VAR condition, 42 kinetochores in >25 cells were measured per experiment. Scale bar, 5 *μ*m. Scale bar zoom-ins, 1 *μ*m. **I)** Hierarchical cluster analysis of Z-score normalized features extracted from data in (B-H) as depicted in Sup. Figure 5C. **J, K)** Plots showing the relation between two SAC proteins on kinetochores with different microtubule-occupancy states. Cells were treated to acquire a mixture of microtubule-occupancy states including the FULL and NULL conditions. Shown here are background (BG)-corrected levels of SAC proteins, plotted against corresponding BG-corrected mCherry levels. At least 72 kinetochores in >30 cells were measured. n = 2, representative experiments are shown here.

In summary, our results indicate that the behaviour of SAC proteins is not uniform on attached kinetochores: all SAC proteins respond to low occupancy states but some are insensitive to further increases in occupancy.

### Immature k-fibers are sufficient to silence the SAC

Our results thus far suggest that at least a substantial part of the SAC machinery is removed from kinetochores well below full occupancy. However, since there is substantial variation in microtubule intensities of fully attached metaphase kinetochores, we next wished to experimentally tweak average maximal occupancy to verify these observations and examine if SAC protein removal at low occupancy is sufficient to silence the SAC. HEC1 is the major microtubule-binding protein on the kinetochore (DeLuca et al., 2004, 2002; Liu et al., 2006; McCleland et al., 2004, 2003; Wigge and Kilmartin, 2001) and its affinity for microtubules is controlled by phosphorylation of its N-terminal tail (Cheeseman et al., 2006; Wei et al., 2007). Designed combinations of phospho-site substitions to phosphomimetic or non-phosphorylatable amino acids (Aspartic acid and Alanine, respectively) generates HEC1 versions with a variety of microtubule-binding affinities, which in cells results in a controlled range of average k-fiber intensities (Zaytsev et al., 2014). We constructed cell lines expressing seven mutant versions of HEC1 to achieve a range of occupancy states (**Sup. Figure 6A-E**). Considering single microtubules can be bound by many HEC1 molecules, our approach enabled creation of uniform HEC1 lawns with specified microtubule-binding affinities, unlike for example diminishing the total amount of HEC1 on kinetochores or mixing high and low affinity HEC1 species. Moreover, the HEC1 mutants simulate the phosphorylation states of kinetochores during unperturbed mitosis (Zaytsev et al., 2014), providing insight into the SAC response during k-fiber maturation.

Cells expressing the HEC1 variants were analyzed for ability to silencing the SAC by time lapse imaging: Occupancy states that cannot silence the SAC are predicted to delay mitosis indefinitely, while those that can, should allow progression. As shown in **Figure 3A and B** and reported before (Etemad et al., 2015; Guimaraes et al., 2008; Zaytsev et al., 2014), cells expressing wild-type HEC1 or HEC1-9A (high microtubule affinity) were able to silence the SAC, whereas those expressing HEC1-9D (low affinity for microtubules) were not. While HEC1-5D was likewise unable to silence the SAC, HEC1-4D (intermediate microtubule affinity) was proficient in SAC silencing, albeit relatively inefficiently (**Figure 3A, B**). Quantitative immunofluorescence showed that HEC1-4D k-fibers were on average 45% of average maximal intensity of HEC1-WT cells, in line with a previous report (Zaytsev et al., 2014), and those of HEC1-5D were ~20% (**Figure 3C, D; Sup. Figure 6**). Simultaneous live imaging of mCherry-MAD2 and eGFP-HEC1-4D showed that HEC1-4D kinetochores had shed most or all of the MAD2 by 30 minutes following mitotic entry (**Figure 3E**). Some kinetochores however had retained substantial MAD2 levels after 93 minutes, likely explaining why mitotic exit was relatively inefficient in these cells. Quantitative immuno-imaging of single attached kinetochores showed that 82% of HEC1-4D kinetochores had MAD2 levels that were as low as those of HEC1-WT kinetochores (**Figure 3F, G**). For comparison, this was true for 30% and 0% of HEC1-5D and HEC1-9D kinetochores, respectively. These data support the hypothesis that kinetochores can inactivate SAC signalling at intermediate (~50%) occupancy states and that SAC silencing becomes more efficient with increasing occupancy.

**Figure 3.**
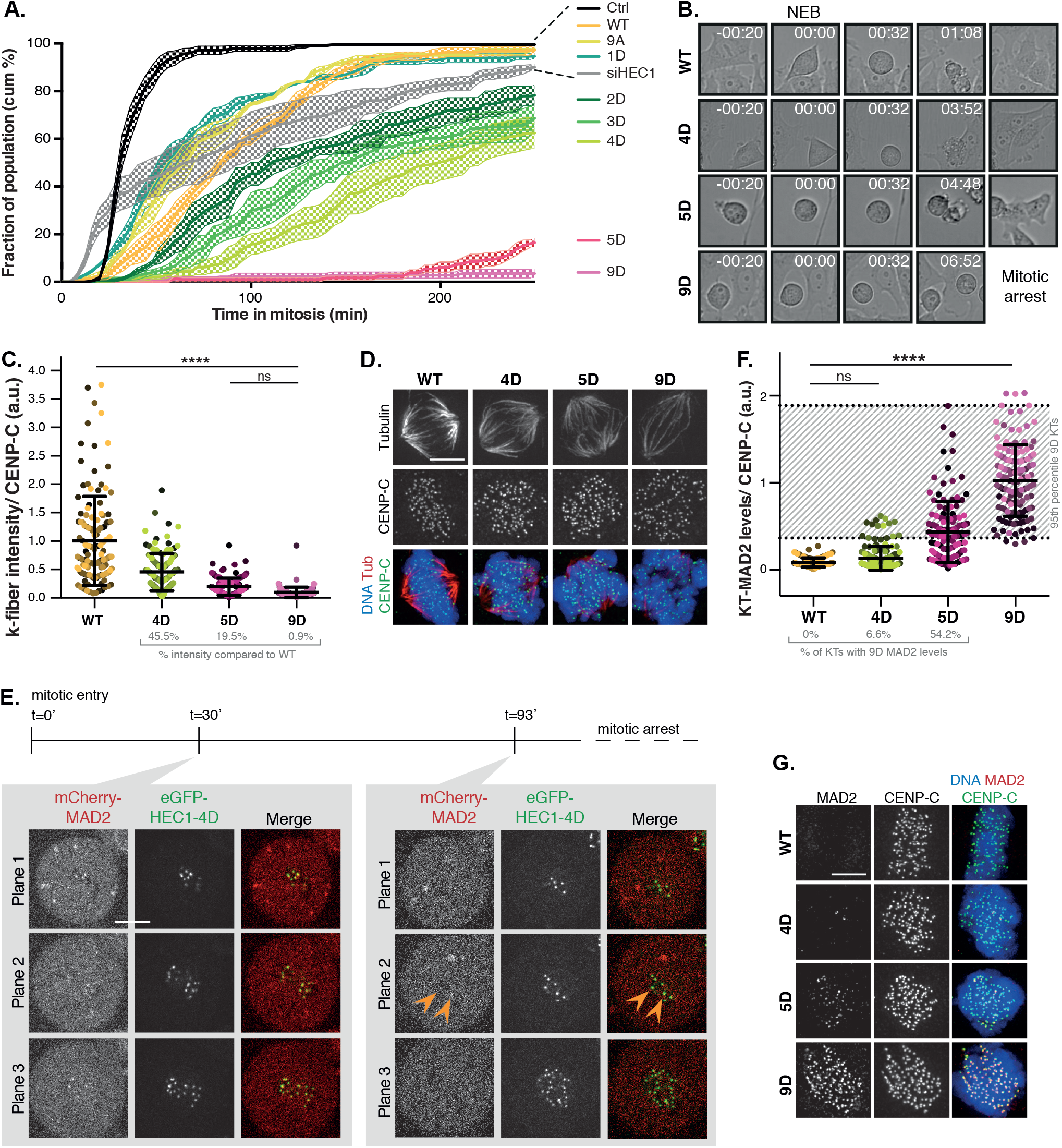
Immature k-fibers are sufficient to silence the SAC. **A, B)** Quantification (A) and representative stills (B) of unperturbed mitotic duration in cells expressing variants of HEC1. At least 50 cells were scored per condition in three independent experiments. Shown here are the average of three experiments (continuous line) and SEM (dotted area). **C, D)** Quantification (C) and representative images (D) of the intensity of k-fibers in cells expressing indicated HEC1 variants. Cells were arrested in metaphase for two hours prior to fixation to allow maximum microtubule occupancy. Each datapoint represents one kinetochore or k-fiber. >118 kinetochores of >5 cells were scored per cell line and different shades of the same color indicate different cells. Channel colors of merged images match those of the labels. n = 2, representative experiment is shown here. For statistical analysis, a one-way Anova test was performed. Scale bar, 5 *μ*m. ****P < 0.0001. **E)** Representative stills of cell in mitotis simultaneously expressing mCherry-MAD2 from its endogenous locus and eGFP-HEC1-4D. Cells were released from a G2/M-block to track mitotic entry and followed ~100 min. Shown here are selected planes (3/16) at two time points. Arrowheads indicate two MAD2-positive kinetochores at t=93’, MAD2 was undetecable on other kinetochores. **F, G)** Quantification (F) and representative images (G) of the intensity of kinetochore(KT)-MAD2 in cells expressing indicated HEC1 variants. Cells were arrested in metaphase for two hours prior to fixation to allow maximum microtubule occupancy. Each datapoint represents one kinetochore or k-fiber. >118 kinetochores of >5 cells were scored per cell line and different shades of the same color indicate different cells. Channel colors of merged images match those of the labels. n = 2, representative experiment is shown here. For statistical analysis, a one-way Anova test was performed. Scale bar, 5 *μ*m. ****P < 0.0001.

## Discussion

Each human kinetochore consists of hundreds of microtubule-binding complexes that each can recruit SAC proteins. In metaphase these kinetochores are bound by ~20 microtubules and have shut down SAC signaling (DeLuca et al., 2005; Guimaraes et al., 2008; McEwen et al., 2001; Wendell et al., 1993). Kinetochores are unlikely to transition from zero to a full complement of microtubules in a single step, yet there is little knowledge about SAC responses to intermediate microtubule occupancies. We show here that key SAC proteins are substantially depleted from kinetochores at ~30% occupancy and are nearly undetectable at ~50% occupancy or above. Our quantitative immuno-imaging of SAC protein levels in relation to microtubule intensities on single kinetochores distinguished two response types. Levels of ZW10, Spindly, MAD1 and MAD2 anti-correlated to microtubule intensities and became not or barely detectable at ~50% occupancy. BUBR1, BUB1 and KNL1-pT180, however, although also declining strongly at low occupancy, were not sensitive to further increases in occupancy and showed variable levels. The behaviors of the members of these two groups are consistent with their mutual physical interactions, and correlate with distinct delocalization mechanisms proposed for these groups. Removal of the BUB group (KNL1-pT180 and by extension BUB1, and BUBR1) requires dephosphorylation by PP1 and decreased localization and activity of MPS1 (Etemad and Kops, 2016; Hiruma et al., 2015; Ji et al., 2015; Nijenhuis et al., 2014). Removal of the RZZ-MAD group occurs through dynein motor activity (Caldas et al., 2015; Kim et al., 2012; London and Biggins, 2014; Matson and Stukenberg, 2014; Silió et al., 2015). Interestingly, MAD1 also interacts with BUB1 in an MPS1-dependent manner (Caldas et al., 2015; Silió et al., 2015), which might explain why its response among all members of the RZZ-MAD group is most similar to that of members of the BUB group. Although MAD1 and MAD2 form a heterotetramer, their behavior in our analyses is not entirely overlapping. The molecualr basis for this is unknown, but MAD2-independent functions for MAD1 at kinetochores have been reported (Akera et al., 2015; Emre et al., 2011).

Cells in which kinetochores reach ~45% occupancy on average (HEC1-4D) can silence the SAC and exit mitosis, while those with ~20% (HEC1-5D) cannot. These data show that a full complement of microtubules such as seen on metaphase kinetochores is not required for SAC silencing. The kinetochore, therefore, acts as a single unit with respect to SAC signaling: when a treshold of bound microtubules is reached, the entire unit switches off its signaling output. This has important implications for our understanding of the SAC as it suggests that the signal from those hundreds of microtubule-binding complexes is quenched by only a few (~7-10) microtubules. We envision several ways in which this can be achieved. First, a few microtubules may be sufficient to pull a stiff kinetochore away from a SAC activating signal (e.g. Aurora B) from inner centromere/kinetochore (Burke and Stukenberg, 2008; Santaguida and Musacchio, 2009; Saurin et al., 2011; Stukenberg and Burke, 2015). We do not favor this hypothesis, as we and others recently showed that distance between sister kinetochores or between inner- and outer kinetochore is not required for SAC silencing (Etemad et al., 2015; Magidson et al., 2016; O’Connell et al., 2008; Tauchman et al., 2015). Second, a low number of microtubules may suffice to elicit a signal that sweeps the kinetochore. For example, phosphatases such as PP1 could be ‘unleashed’ from a site of recruitment/activation upon a threshold of microtubule binding. Concurrent with sufficient MPS1 displacement, this could switch the SAC signal to an OFF state. It is unclear, however, how dynein-mediated removal of RZZS-MAD proteins would occur in such a scenario. Third, the kinetochore may be flexible, allowing only a few microtubules to engage the majority of microtubule-binding complexes and thus displace sufficient MPS1 molecules and recruit sufficient PP1 and dynein molecules to achieve substantial SAC protein delocalization. Transition to full occupancy may then be facilitated by kinetochore flexibility and many low affinity microtubule interactions (Etemad and Kops, 2016). Fourth, attachments may be highly dynamic, engaging and disengaging kinetochores frequently. This may allow most of the microtubule-binding complexes to briefly bind microtubules and shed SAC proteins. A sufficiently high frequency of these labile interactions could conceivably render the kinetochore in a SAC silenced state.

## Materials and Methods

### Cell Culture and transfection

HeLa and HeLa FlpIn cells were grown in DMEM (Sigma; 4,5 g glucose/L) supplemented with 8% tetracycline-free FBS (Bodingo), penicillin/streptomycin (Sigma; 50 μg/ml), GlutaMAX (Gibco; 5 mL), and hygromycin (200 μg/ml) or puromycin (1.6 μg/ml). Plasmids were transfected using Fugene HD (Roche) according to the manufacturer’s instructions. To generate stably integrated constructs, HeLa FlpIn cell lines were transfected with pCDNA5-constructs and pOG44 recombinase simultaneously in a 1:9 ratio(Klebig et al., 2009). Constructs were expressed by addition of 1 μg/ml doxycycline for 24h. siHEC1 (custom; Thermo Fisher Scientific; 5’-CCCUGGGUCGUGUCAGGAA-3’) and siGAPDH (Thermo Fisher Scientific; D-001830-01-50) was transfected using HiPerfect (Qiagen) according to manufacturer’s instructions.

### Plasmids

pCDNA5-pEGFP-HEC1 constructs and cloning strategies are described in (Nijenhuis et al., 2013). Other constructs were made using site-directed mutagenesis by PCR.

### CRISPR/Cas9 genome editing of *MAD2* and *BUB1* loci

Inserting the gene for mCherry into the endogenous loci of *MAD2* and *BUB1* was performed using self cloning CRISPr strategy (Arbab et al., 2015). In brief: 3xFLAG-spCas9 was subcloned from spCas9-BLAST to pcDNA3-MCS-IRES-PURO using NdeI/EcoRI restriction digestion to allow selection for spCas9 expression in HeLa FLPin cells. To generate HA-mCherry, pcDNA3Zeo-CyclinB-mCherry (Kuijt et al., 2014) was used as template and amplified by PCR using forward (AAGCTTTACCCGTACGACGTGCCAGATTACGCTGTGAGCAAGGGCGAGGAGG) and reverse (gcgccgTCTAGATCCGCAGCCACCGCCAGATCCGCCCTTGTACAGCTCGTCCATGC) primers. The PCR fragment was digested with HinDIII/XbaI and ligated into pcDNA3.0 via HinDIII/XbaI.

To create the homology arms, three consecutive PCR-amplifications were done on HA-mCherry template to create MAD2 (120bp 5’ of ATG and 120bp 3’) or BUB1 (119bp 5’ of ATG and 124bp 3’). spCas9 was directed to MAD2 using sgRNA: G-AGCTGCAGCGCCATGGCC, or BUB1 using G-TCCTCTGGCCATGGACACCC. Both homology arms and sgRNA template were subcloned into pJet1.2 (ThermoFischer) and sequence verified before PCR fragments were generated for transfection.

To generate HeLa FlpIn cells with endogenous tagged MAD2 or BUB1 cells were transfected with 1.5μg spCas9-IRES-PURO, 1.5μg sgPAL7-Hygro, 3μg homology PCR template and 3μg sgRNA PCR template at a ratio 1:3 DNA:Lipofectamine LTX (ThermoFischer). 24 hours after transfection, 1 μg/ml puromycin and 200 μg/ml Hygromycin B were used for 48 hours after which cells were grown till confluency in a 10 cm petri dish. HeLa FlpIn cells subsequently FACS-sorted as single cells on using BD FACSAria FUSION (640nm excitation laser, autofluorescence 670nm/30 vs 651nm excitation laser, 610nm/20 mCherry channels, 100μm nozzle, 2.0 flowrate). Clones were verified to have correct labelling of MAD2 or BUB1 by PCR on genomic DNA, western blotting and live cell immunofluorescence microscopy.

### Knockdown and reconstitution experiments

To knockdown and reconstitute HEC1 in HeLa FLpIn cell lines, cells were transfected with 40 nM HEC1 or mock siRNA and arrested in early S phase for 24 h by addition of thymidine (2 mM). Cells were then allowed cell cycle re-entry by washing the cells once with appropriate media. 8 h after thymidine release, cells were treated with doxycycline (1 μg/ml), arrested again using thymidine and incubated with both reagents for 16 h after which they were released from thymidine and further processed. Cells processed for immunofluorescence imaging of SAC proteins and k-fibers were released from thymidine in RO (5 μM) and incubated for 8 h or more. Subsequently, cells were washed three times with warm media, incubated between each wash for 5 minutes at 37 ° C, and incubated for 120 minutes with nocodazole (3,3 μM, ‘NULL’ condition), or MG-132 (5 μM, ‘FULL’ condition). Then, cells were fixed and processed appropriately. To fix cells before all kinetochores had reached full occupancy (the ‘VAR’ condition), cells were fixed and processed 25 min after release from RO. For immunofluorescence imaging of cells expressing HEC1 variants, cells were treated with MG-132 (5 μM) for 120 minutes prior to fixation.

### Live cell imaging

For live cell imaging experiments, cells were plated in 24-well plates (Greiner bio-one), and subjected to DIC microscopy on a Nikon Ti-E motorized microscope equipped with a Zyla 4.2Mpx sCMOS camera (Andor). A 20x 0.45 NA objective lens (Nikon) was used. Cells were kept at 37°C and 5% CO_2_ using a cage incubator and Boldline temperature/CO2 controller (OKO-Lab). Images were acquired every 4 minutes at 2×2 binning and processed by Nikon Imaging Software (NIS). Analysis of live cell imaging experiments was carried out with ImageJ software and time in mitosis was defined as the time between nuclear envelope breakdown and anaphase-onset or cell flattening.

Live cell imaging of mCherry-tagged MAD2 and BUB1 in single cells was performed on a Nikon Time-Lapse system (Applied Precision/GE Healthcare) equipped with a Coolsnap HQ2 CCD camera (Photometrics) and Insight solid-state illumination (Applied Precision/GE Healthcare). Cells were plated in 8-well plates (μ-Slide 8 well, Ibidi) and imaged in a heated chamber (37°C and 5% CO^2^) using a 60×/1.42 NA or 100×/1.4 NA UPlanSApo objective (Olympus) at 2×2 binning. Images were acquired every 15 seconds (for the mCherry-MAD2 cells), or 1 min (for the mCherry-BUB1 cells), and deconvolved using standard settings in SoftWorx (Applied Precision/GE Healthcare) software. Multiple z layers were acquired and projected to a single layer by maximum intensity projection. For simultaneous imaging of GFP(-HEC1) and mCherry-MAD2, the same system was used. Cells were plated in 8-well plates (μ-Slide 8 well, Ibidi), treated with siRNA, thymidine and RO as described above. Images were acquired 30 and 60 minutes after mitotic entry, and then every three minutes.

### Immunofluorescence and image quantification

For fixed cell immunofluorescence microscopy, cells plated on round 12-mm coverslips (No. 1.5) were pre-extracted with 37°C 0.1% Triton X-100 in PEM (100 mM Pipes (pH 6.8), 1 mM MgCl2, and 5 mM EGTA) for ±45 s before fixation (with 4% paraformaldehyde) for 10 min. Coverslips were washed twice with cold PBS and blocked with 3% BSA in PBS for 16 h at 4°C, incubated with primary antibodies for 16 h at 4°C, washed 4 times with PBS containing 0.1% Triton X-100, and incubated with secondary antibodies for an additional hour at room temperature. Coverslips were then washed twice with PBS/0.1% Triton X-100, incubated with DAPI for 2 min, washed again twice with PBS and mounted using Prolong Gold antifade (Invitrogen). For cold-shock experiments, cells were placed on ice water in 500 μL media for 8 minutes prior to pre-extraction and fixation with the appropriate buffers.

All images were acquired on a deconvolution system (DeltaVision Elite; Applied Precision/GE Healthcare) with a 100×/1.40 NA UPlanSApo objective (Olympus) using SoftWorx 6.0 software (Applied Precision/GE Healthcare). Deconvolution is applied to all images and maximum projection is shown in figures, except for panel (A) in **Figure 2**, which is a sum projection image, and panel 3C in which single planes are shown. For quantification of immunostainings, all images of simultaneously stained experiments were acquired with identical illumination settings. For analysis of the HEC1 mutant expressing cell lines, cells expressing comparable levels of exogenous protein were selected for analysis and analyzed using ImageJ. For measurement of protein levels and k-fiber intensities on single kinetochores, kinetochores were selected in max projection images. The 7-8 slices that contained a single kinetochore and corresponding k-fiber were selected and sum projection images were used for quantification. Line plots were used to determine the highest intensity at kinetochores/k-fibers and local background was subtracted from these values (**Sup. Figure 1**). The same method was applied to determine protein levels, k-fiber intensity and CENP-C levels. k-fiber and protein measurements were normalized to CENP-C to correct for biological and technical variation between kinetochores. Further normalization steps include normalization of k-fiber levels to the median of k-fiber levels measured in FULL conditions, and normalization of protein levels to the levels measured for the median of the same protein in the NULL conditions.

### Data analysis

Data analysis was performed in R (3.3.2) using the pheatmap (1.0.8, CRAN), MarkdownReports (2.5, DOI: 10.5281/zenodo.594683) packages. Raw measurement per kinetochore were normalized as described in ‘Immunofluorescence and image quantification’.

To quantify features of individual occupancy response-curves in transient (left) and steady (right) phase separately, piecewise linear regression was applied, where a breakpoint separates the two phases. Each feature is extracted from either the FULL, NULL or the VAR datasets as denoted. These features are: VarianceInPhase2 (variance in protein concentration in the steady phase, right of the split point,), MedianPhase2 (median protein concentration in the steady phase), Null2Full decrease (relative protein decrease between the two conditions defined as the ratio of median protein levels NULL over FULL attachment, corresponds to % values in **Figure 1B**), MedianFull (median protein levels in the full condition), BreakPoints (X or Tubulin-coordinate of the split point in the piece-wise linear regression), and Slope (slope of the fitted line in the transient phase, left of the split point).

To investigate variance across proteins at full attachment, measured values were tested for normality (**Sup. Figure 7**). Based on these results, Levene’s test was used to compare variances. Code availability: Upon acceptance of the manuscript, the source code for the analysis, raw quantification data will be available “as-is” under GNU GPLv3 at https://github.com/vertesy/Kinetochore.

### Immunoblotting

Cells were treated as described above and entered mitosis in the presence of Nocodazole. Cells were collected and lysed in Laemmli lysis buffer (4% SDS, 120 mM Tris (pH 6.8), 20% glycerol). Lysates were processed for SDS-PAGE and transferred to nitrocellulose membranes for immunoblotting. Immunoblotting was performed using standard protocols. Visualization of signals was performed on a scanner (Amersham Imager 600) using enhanced chemiluminescense.

### Antibodies

The following primary antibodies were used for immunofluorescence imaging: CENP-C (guinea pig polyclonal, 1:2,000; Sigma-Aldrich), α-tubulin (mouse monoclonal, 1:10,000; Sigma-Aldrich), HEC1 (mouse monoclonal 9G3, 1:2,000; Abcam), GFP (custom rabbit polyclonal raised against full-length GFP as antigen, 1:10,000 (Jelluma et al., 2008)), GFP (mouse monoclonal, 1:1,000; Roche), MAD2 (custom rabbit polyclonal raised against full-length 6×His-tagged MAD2 as antigen, 1:2,000 (Sliedrecht et al., 2010)), BubR1 (rabbit polyclonal, 1:1,000; Bethyl), BUB1 (rabbit polyclonal, 1:1000, Bethyl), Spindly (rabbit polyclonal, 1:1000, Bethyl), ZW10 (rabbit polyclonal, 1:1000, Abcam), MAD1 (rabbit polyclonal, 1:1000, Santa Cruz), RFP (rat monoclonal, 1:1000, Chromotek) GFP-Booster (Atto 488, 1:500, Chromotek). Secondary antibodies were highly crossed absorbed goat anti-guinea pig Alexa Fluor 647, anti-rat Alexa Fluor 568, and goat anti–rabbit and –mouse Alexa Fluor 488 and 568 (Molecular Probes).

## Acknowledgements

We thank Jennifer DeLuca for reagents, members of the Kops lab for discussions, and Sophie Dumont and Jonathan Kuhn for sharing unpublished data. This study was supported by Oncode institute, which is partly funded by the Dutch Cancer Society, and by the Netherlands Organisation for Scientific Research (NWO-Vici 865.12.004).

---

**Supplemental Figure 1.**
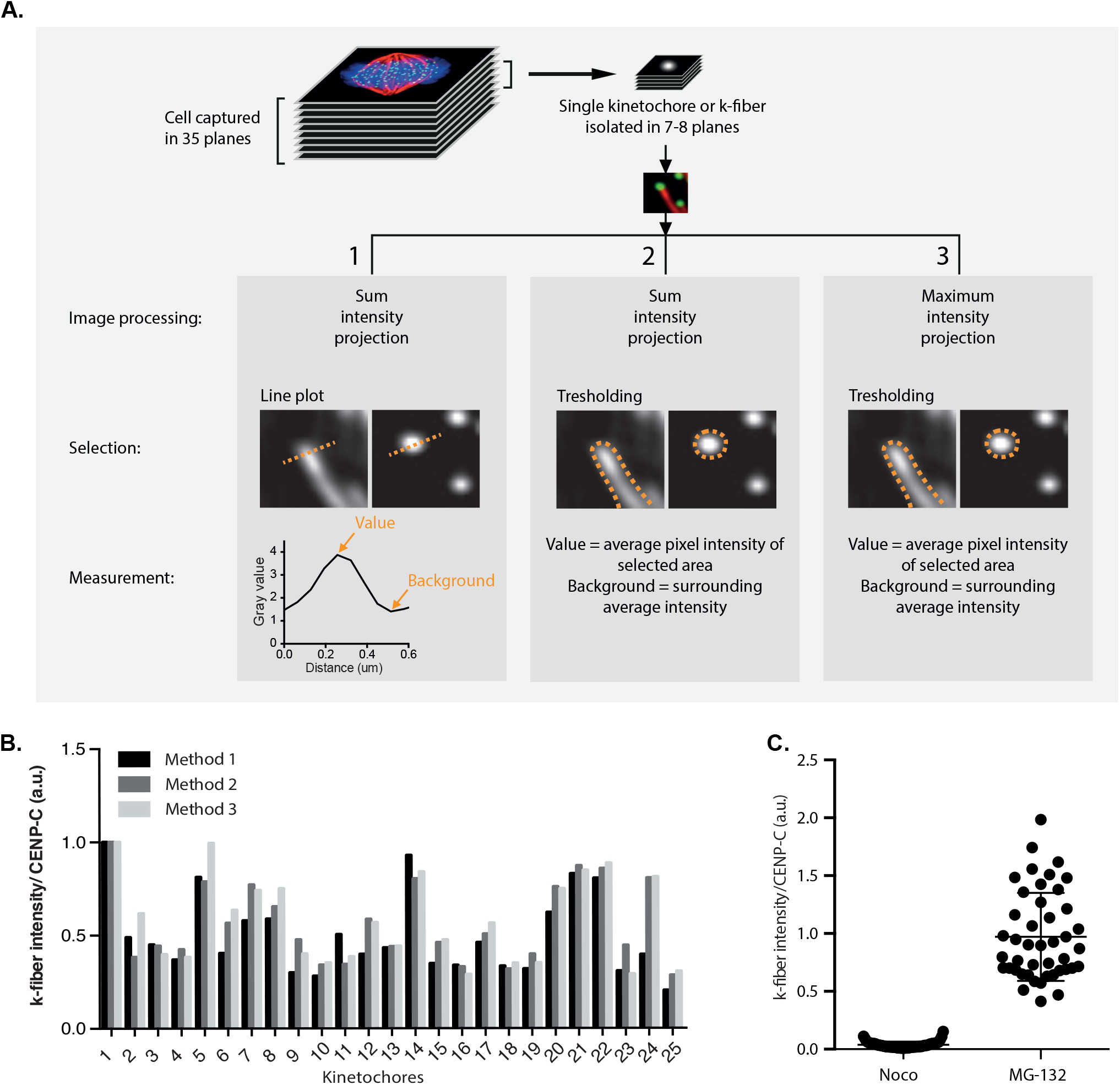
Summary and validation of methods used to measure k-fiber intensity. **A)** Several image-processing (sum or maximum intensity projections), selection and measuring methods (line plots or thresholding and corresponding measurements) were combined into three pipelines and applied to the same images. **B)** Bar graph showing intensity of individual k-fibers normalized to CENP-C levels. Each k-fiber is measured multiple times using the methods depicted in (A). Data is normalized to the values measured for k-fiber 1 and is compiled from 10 cells. **C)** k-fiber intensity over CENP-C is plotted for cells treated with nocodazole or MG-132 after entry into mitosis for a duration of 1,5 hours. Each data point represents one k-fiber. All data points are normalized to the average K-fiber intensity measured in MG-132-treated cells.

**Supplemental Figure 2.**
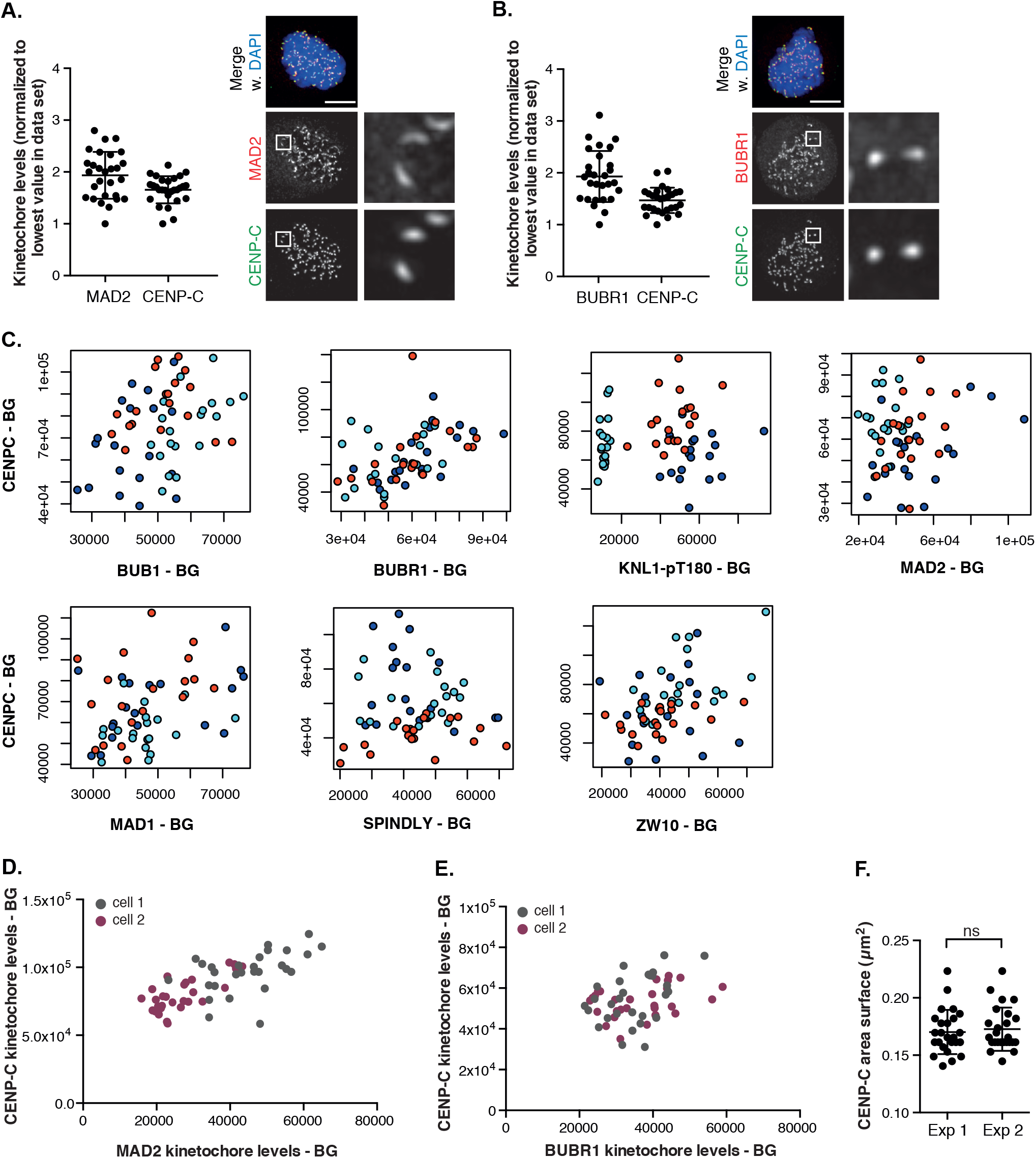
High variability of protein levels on unattached kinetochores is independent of CENP-C levels, the kinetochore size, antibody penetration or differences between replicates. **A, B)** Kinetochore levels of SAC proteins is highly variable within the same cell contrary to the levels of CENP-C measured on the same kinetochores. Figures show immunofluorescence images and quantification of protein levels at kinetochores of a nocodazole-treated cell. Each data point represents one kinetochore. Levels of each protein are normalized to the lowest value in the data set to allow comparison of variance between proteins. Scale bar, 5 *μ*m. **C)** Staining efficiency is similar in the biological replicates of experiments shown in Figure 1. Graphs show background (BG)-corrected levels of SAC proteins at unattached kinetochores, plotted against corresponding BG-corrected CENP-C levels. Different colors depict individual experiments. **D, E)** Cell-to-cell variability due to antibody penetrance is corrected through normalization against CENP-C in all experiments. To exclude variance in the data caused by differences in antibody penetration proficiency, background (BG)-corrected levels of SAC proteins at unattached kinetochores of single cells are plotted against corresponding BG-corrected CENP-C levels. For each SAC protein, the kinetochore levels of two simultaneously handled cells are shown. **F)** Surface area of CENP-C in two independent experiments. Kinetochores of 5 cells are used per experiment and each data point represents one kinetochore. For statistical analysis, an unpaired Student’s t-test was performed.

**Supplemental Figure 3.**
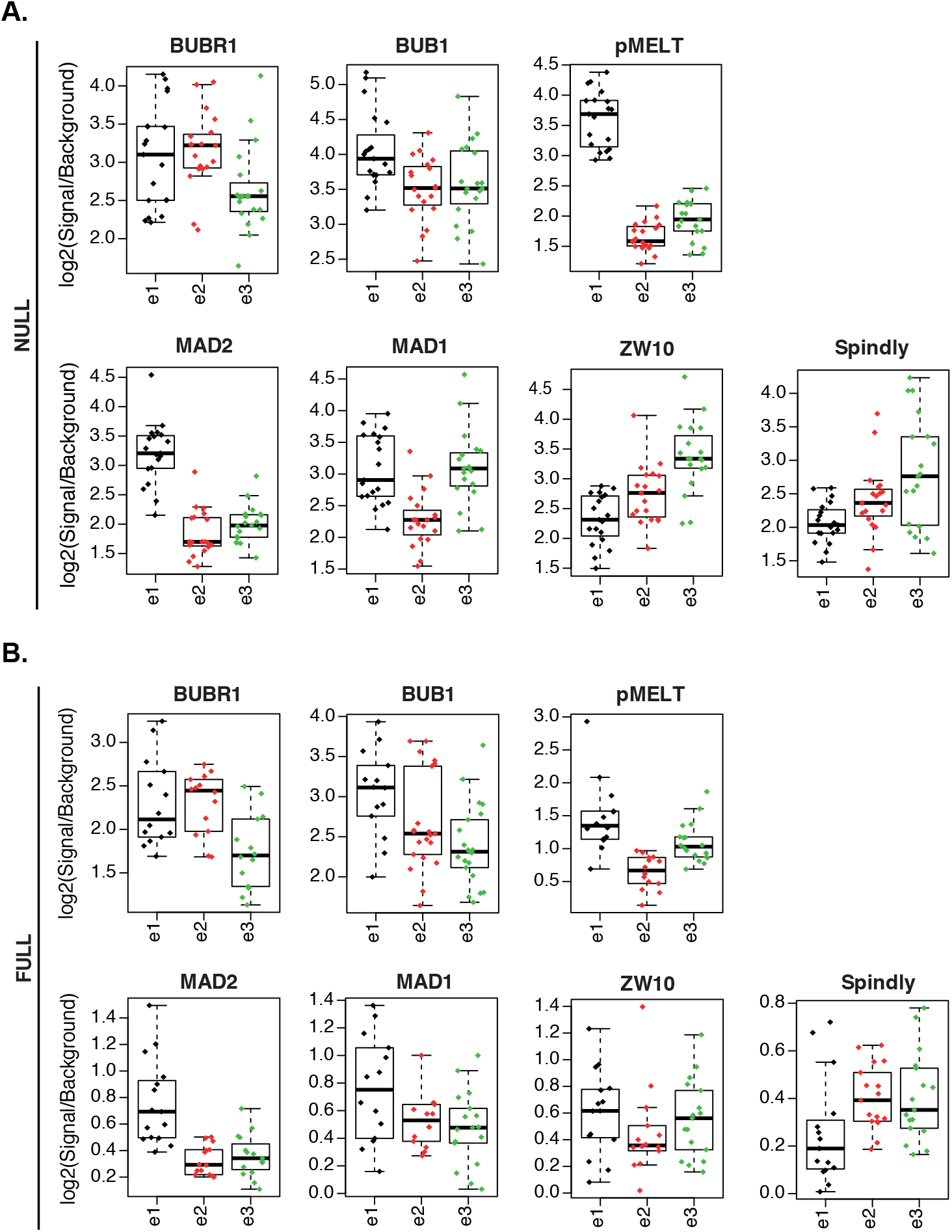
Variability of protein occupancy in FULL condition is not due to signal to noise ratio. **A, B)** Signal-to-noise ratios of SAC proteins measured on kinetochores of unattached (A) and fully microtubule-occupied kinetochores (B). Each biological replicate is plotted separately and represented by ‘e1’, ‘e2’, and ‘e3’. All proteins and experiments in NULL condition show similar signal-to-noise ratios, suggesting that variance is equally affected by noise across all experiments. In FULL condition signal-to-noise ratio is much better in proteins with high variation (BUB1, BUBR1, and pMELT), strongly supporting that the observed behavior in Figure 1A-D is not due to skewed signal intensities.

**Supplemental Figure 4.**
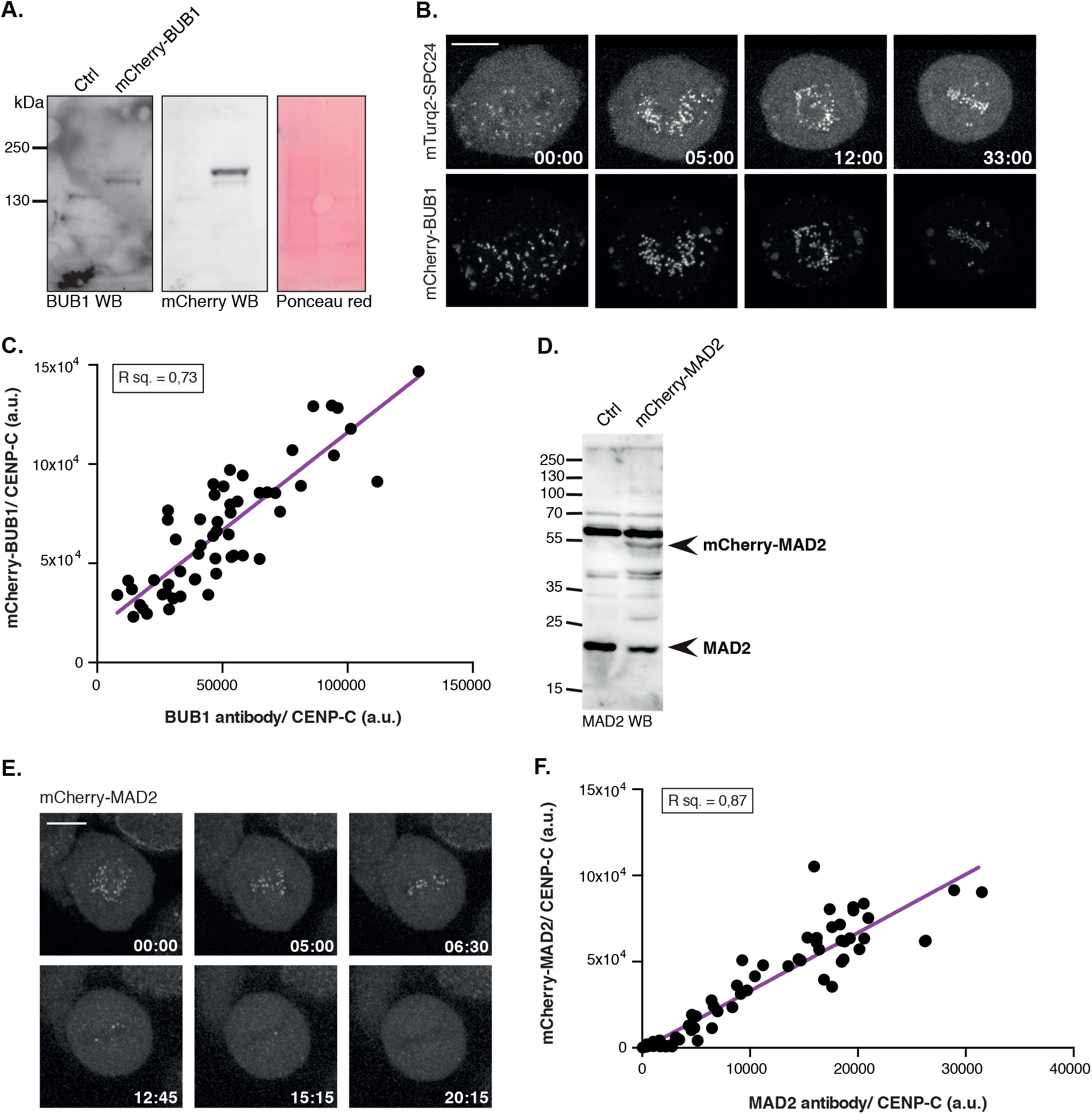
Visualization of SAC proteins using antibodies is similar to protein visualization using a fused fluorescent tag expressed from the endogenous locus of the same protein. **A)** Western blot of lysates of unsynchronized control versus mCherry-BUB1 expressing cells. **B)** Stills of representative movie of a cell expressing mCherry-BUB1 and mTurq2-SPC24 simultaneously. mTurq2-SPC24 is shown as a reference. Stills are maximum projection images. Scale bar, 10 *μ*m. **C)** Scatter plot showing background (BG)-corrected levels of BUB1 visualized by a polyclonal antibody, plotted against corresponding BG-corrected levels of mCherry. Cells were fixed in prometaphase to create a range of attachments. 55 kinetochores of >10 cells were measured for this experiment. **D)** Western blot of lysates of unsynchronized control versus mCherry-MAD2 expressing cells. **E)** Stills of movie following a mCherry-MAD2 expressing line through mitosis. Stills are maximum projection and bleach-corrected images. Scale bar, 10 *μ*m. **F)** Scatter plot showing background (BG)-corrected levels of MAD2 visualized by a polyclonal antibody, plotted against corresponding BG-corrected levels of mCherry. Cells were fixed in prometaphase to create a range of attachments. 72 kinetochores of >20 cells were measured for this experiment.

**Supplemental Figure 5.**
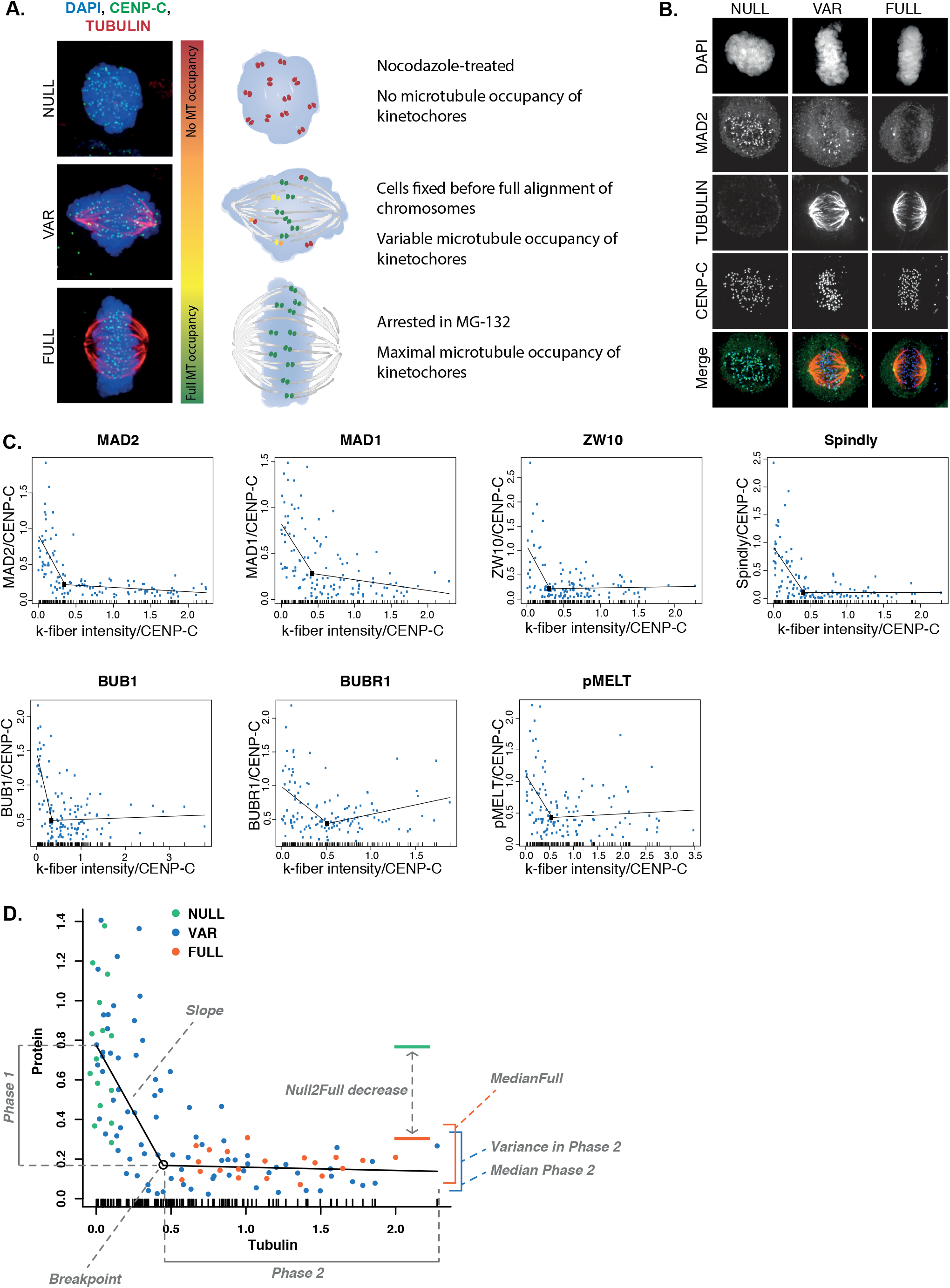
Creating and measuring a mixture of kinetochore-microtubule attachment states and their corresponding SAC protein levels. **A, B)** Graphic depicting the conditions used to create a mixture of attachment states (A), and representative immunofluorescent images (B) to illustrate visualization of k-fibers and SAC proteins when NULL, VAR and FULL conditions are applied. Scale bars, 5 *μ*m. **C)** Graphs depicting curve fits performed on the data shown in Figure 2 B-H. After piecewise linear regression, a variety of features as depicted in (D) was extracted from the fit and the data to describe the relation of tubulin to protein of interest. **D)** Illustration depicting the features of SAC protein localization in relation to microtubule occupancy. Features are: variance of data along the Y-axis at the stationary stage (after breakpoint, VariancePhase2), rate of decrease in protein levels to minimum levels (before breakpoint, Slopes), tubulin level at which proteins reach their minimum levels (Breakpoints), median of protein level at stationary stage (MedianPhase2), median protein level in the FULL condition (MedianFull), and the decrease of FULL data set in relation to the NULL data (Null2Full).

**Supplemental Figure 6.**
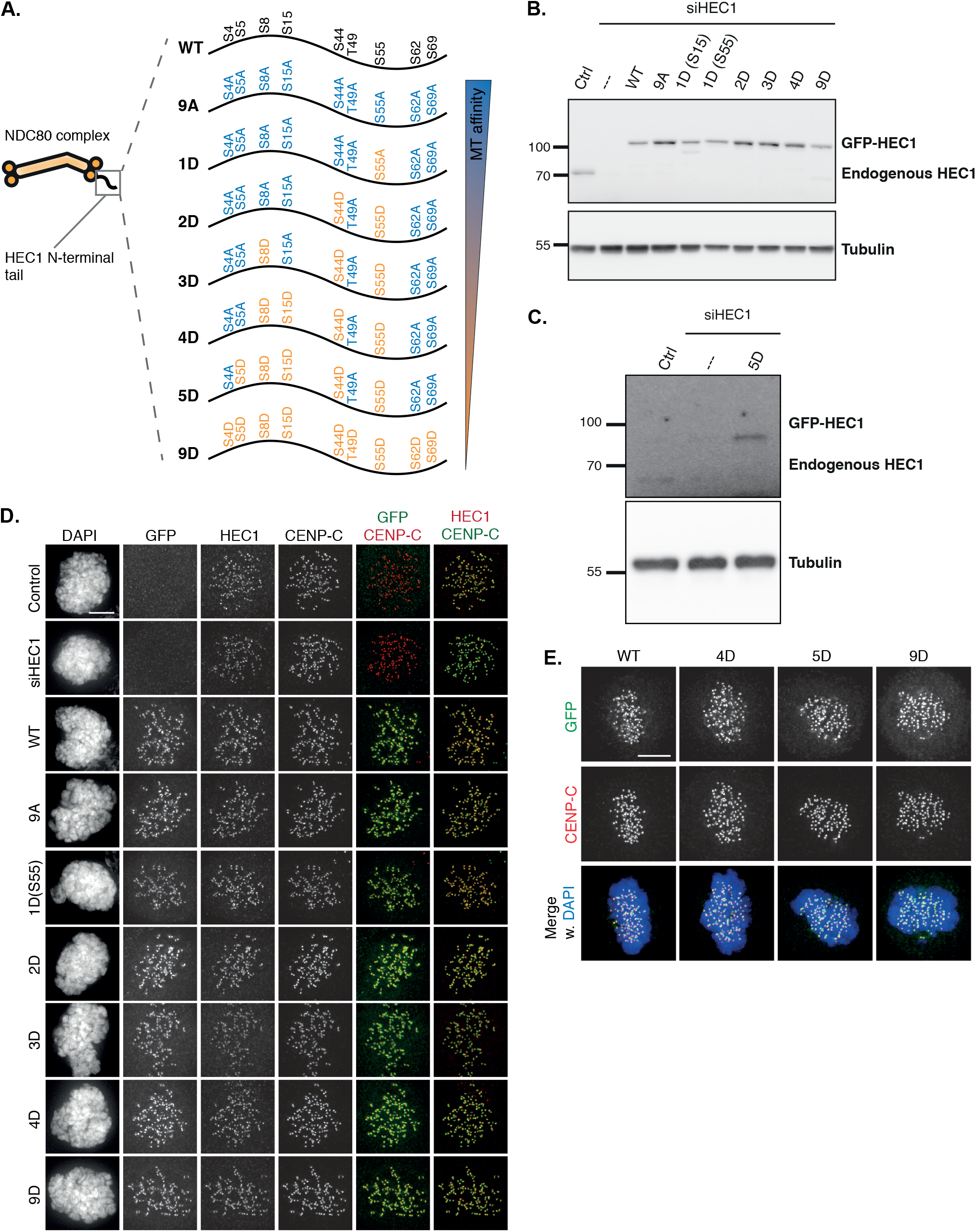
Characterization of cells expressing variants of HEC1. **A)** Overview of the mutants used in this paper. The HEC1 variants have phosphomimetic or phosphodead aminoacid subtitutions in all nine known phosphorylation sites in the N-terminal tail of HEC1. **B, C)** Representative immunoblots showing HEC1 knock down and doxycycline-induced expression of siRNA-resistant eGFP-tagged versions of HEC1 in a mitotic population. As both HEC1-1D versions behave similarly (Etemad et al., 2015; Zaytsev et al., 2015, 2014), further experiments were performed using HEC1-1D(S55D). **D, E)** Representative immunofluorescent images of indicated proteins in nocodazole- (D) or metaphase- (E) arrested cells. Channel colors of merged images match those of the labels. Scale bars, 5*μ*m.

**Supplemental Figure 7.**
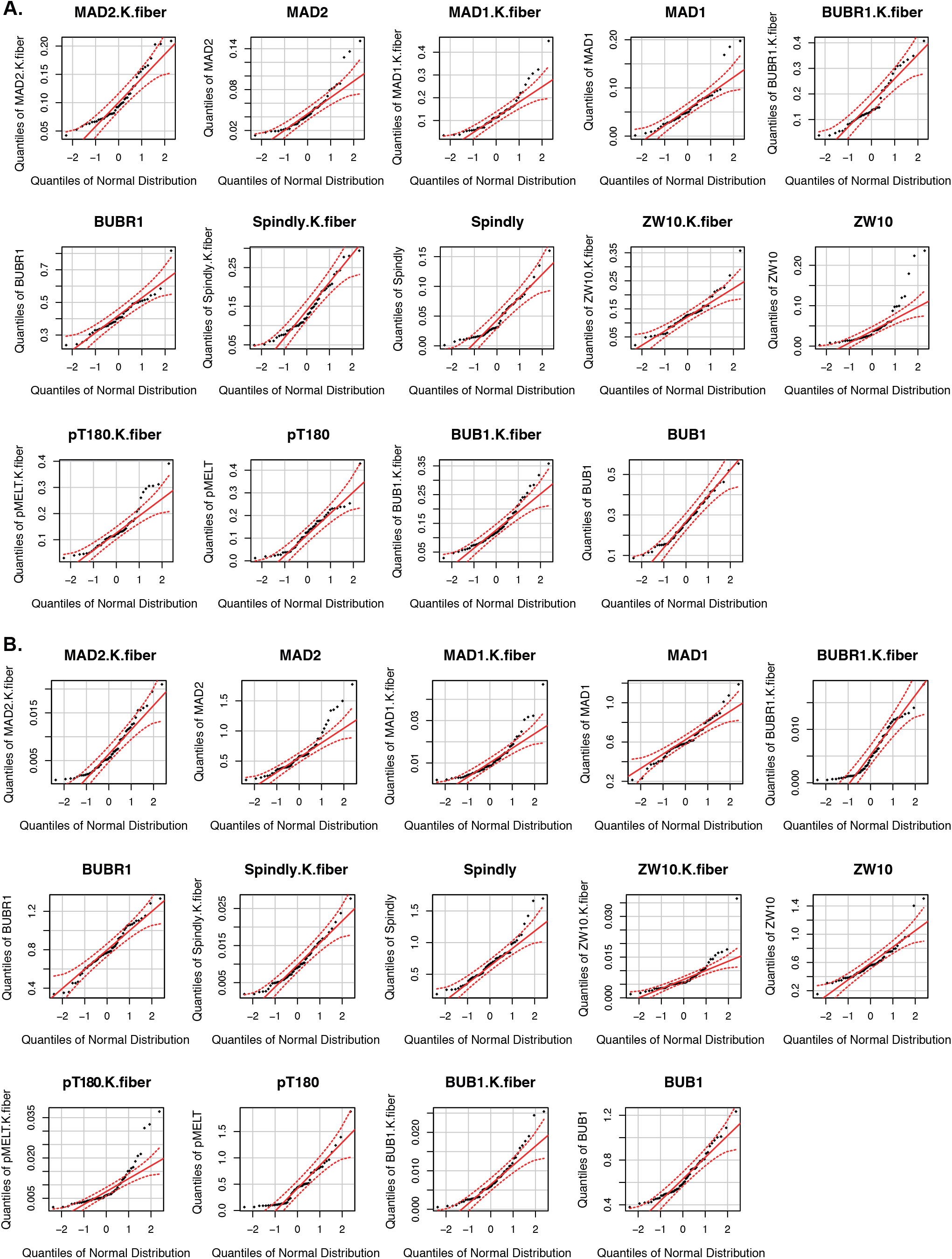
Measured intensities are not normally distributed. **A, B)** QQ plots displaying quantiles of datasets as a function of the quantiles of a normal distribution for the FULL (A) and NULL (B) data sets.

